# Simulation and Analysis of Animal Movement Paths using Numerus Model Builder

**DOI:** 10.1101/2019.12.15.876987

**Authors:** Wayne M. Getz, Ludovica Luisa Vissat, Richard Salter

## Abstract

Animal movement paths are represented by point-location time series called relocation data. How well such paths can be simulated, when the rules governing movement depend on the internal state of individuals and environmental factors (both local and, when memory is involved, global) remains an open question. To answer this, we formulate and test models able to capture the essential statistics of multiphase versions of such paths (viz., movement-phase-specific step-length and turning-angle means, variances, auto-correlation, and cross correlation values), as well as broad scale movement patterns. The latter may include patchy coverage of the landscape, as well as the existence of home-range boundaries and gravitational centers-of-movement (e.g., centered around nests). Here we present a Numerus Model Builder implementation of two kinds of models: a high-frequency, multi-mode, biased, correlated random walk designed to simulate real movement data at a scale that permits simulation and identification of path segments that range from minutes to days; and a model that uses statistics extracted from relocation data—either empirical or simulated—to construct movement modes and phases at subhourly to daily scales. We evaluate how well our derived statistical movement model captures patterns produced by our more detailed simulation model as a way to evaluate how well derived statistical movement models may capture patterns occurring in empirical data.

## 1 INTRODUCTION

The purpose of this paper is to present and evaluate the utility and performance of a new method for numerically simulating an animal movement path using a model derived from empirically acquired, relatively high-frequency, relocation data (i.e., minute-by-minute sequential locations of an individual as it moves over a landscape). Following (Getz 2019), we refer to this model as an M^3^ (M-cubed) model; which is an acronym, clarified below, for <u>m</u>ulti-CAM <u>m</u>etaFuME <u>M</u>arkov model. High frequency relocation data appropriate for the extraction of an M^3^ model is relatively expensive to come by and not yet widely shared by groups currently collecting such data for specific systems. Here, in place of high-frequency empirical data, we use relocation data generated from a multi-mode, biased, correlated, random walk (MBCRW) model. The multi-mode component is used to create specific kinds of local movement patterns (e.g., browsing within and movement among resource acquisition patches). The biases component is used to add directed movement to the global path structure. We evaluate the extent to which an M^3^ model is able to identify different local modes in a movement path, as induced by step-length and turning-angle switches among modes in the movement patterns produced by our MBCRW model. We also discuss how the M^3^ method for building an extracted model from high-frequency movement data might be enhanced to enable it to capture global structures in animal movement paths.

The standard form of a relocation data set in the *x-y* plane for an individual movement path that nominally begins at *t* = 0 and ends at *t* = *t_f_*, takes the form

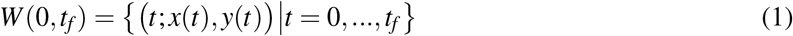

For convenience, we selected the units of *t* to be the inter relocation point interval. Such movement paths reflect elements of the underlying movement behavior repertoires of animals that typically are influenced by seasonally related episodic and cyclic phases. Embedded within the structure of seasonal phases we are likely to observe finer movement modes that switch amongst one other at a within-day scale. Thus movement at the daily scale can be viewed as a mixture of movement behavioral modes, also known as canonical activity modes (CAM; Getz and Saltz 2008), which themselves can be segmented into fixed-length segments of fundamental movement elements (FuMEs; Getz and Saltz 2008) that are referred to as metaFuME segments. Additional factors influencing the movement pathway structure of individuals may include an individual’s physiological or emotional state, the time-of-day, territoriality, distances from attractive centers (e.g., watering points, nests), the presence of both conspecifics and heterospecific competitors and enemies, the physical and ecological structure of the landscape itself, and memories of favorable locations both near and far that provide access to food, water, and safety.

Our ability to extract the role that the various factors listed above play in determining the mode and phase structure of *W*(0, *t_f_*) (Eq. 1) depends on the resolution of the data—specifically, the frequency at which it was collected or the units of *t*. Coarse data (e.g., hourly or subdaily) may only permit us to uncover environmental factors driving seasonal movement phases, intermediate resolution data (e.g., subhourly) may only allow us to uncover environmental and internal state factors driving diel (daily) cycles, while high resolution data (minute or sub-minute) is needed to answer questions that relate to behavior lasting a few to several minutes.

No matter the resolution of the relocation data, however, the first step in dealing with such data is to extract bivariate step-length (SL, *s*(*t*)) and turning-angle (TA, *a*(*t*)) time-series, using the approach described below. Once this has been done, then various methods have been used to extract information from the derived bivariate time series (*s*(*t*), *a*(*t*), along with available covariate data (environmental, individual state, location of conspecifics, and also of heterospecific competitors and enemies), to confront the challenge mentioned above. The efficacy of such methods—which include hidden Markov models (Zucchini et al. 2016) and behavioral change point analysis (Gurarie et al. 2009, Gurarie et al. 2016)—is most easily evaluated using data that is derived from processes known to be causative in driving the observed movement patterns. One way to do this is to analyse simulation data derived from movement models of varying complexity (e.g., MCBRW models) to see how much information can be extract from the relocation and covariate data sets.

In summary, relocation data sets of the type depicted in Eq. 1 can either be empirical, (which we flag using a hat notation—i.e., *Ŵ*(0, *t_f_*)) or derived from simulation models (which we flag using a hat notation— i.e., 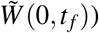. In addition, the simulation model can be fitted to empirical relocation data using some appropriately formulated “best-fitting” procedure.

We use the notation 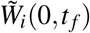 to denote the *i^th^* instance of the relocation data set derived from the *i^th^* simulation of a model that “best fits” an empirical relocation data set Ŵ(0, *t_f_*). In addition, we use calligraphic notation to denote sets of sets, so that an ensemble of *N* simulated “best fitting” walks 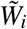 is denoted by

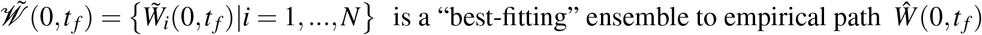

The challenge lies in first finding a suitable class of appropriate complexity models with which to model the process producing the data *Ŵ*(0, *t_f_*). It then moves on to finding the best-fitting set of parameters for this model such that the data *Ŵ*(0, *t_f_*) are more likely to be lie within the ensemble 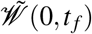 than any other ensemble that can be generated by the model using a different set of parameters. Solving the above problem represents a program of research and study that will be with us for decades to come because it is both model-structure and estimation-method dependent. Models with improved and refined structures will be proposed over time as we learn more about the process that generate the movement paths of individuals belonging to particular species. In addition, we will steadily develop better methods of estimation as computational technologies and parameter-fitting algorithms improve over time.

The approach we take draws upon a hierarchical, appropriate complexity, segmentation framework (Getz 2019) that can be used to breakdown empirical life-history tracks (LiTTs) into a series of diel activity routines (DARs) that belong to particular life history phases (LiMPs) of individuals’ LiTTs (e.g., a ranging phase versus a migratory phase—see Getz 2019 for further discussion). These DARs can then be further segmented into sequences of different types of canonical activity modes (CAMs; Fig. 1A.) that, in turn, can be modeled as sequences of “meta” fundamental movement elements (metaFuMEs). Additionally, we assume that each type of CAM is constructed from the same basis set of metaFuMEs, with differences among CAM types characterized by the lengths and frequencies in the occurrence of particular metaFuME sequences (Fig. 1B.). As an aside, we note that fundamental movement elements themselves, as discussed in (Getz and Saltz 2008, Getz 2019), cannot generally be extracted directly from relocation data. Rather, current data are typically only suitable for identifying CAMs from step-size and turning-angle time series using hidden Markov model (HMM) and behavioral change point analysis (BCPA) methods. Such methods have been used to identify CAMs at a particular sampling frequency resolution. Aggregating all the CAMs of the same type together allows us to construct bivariate step-length and turning-angle distributions for the CAM ensembles.

**Figure 1:**
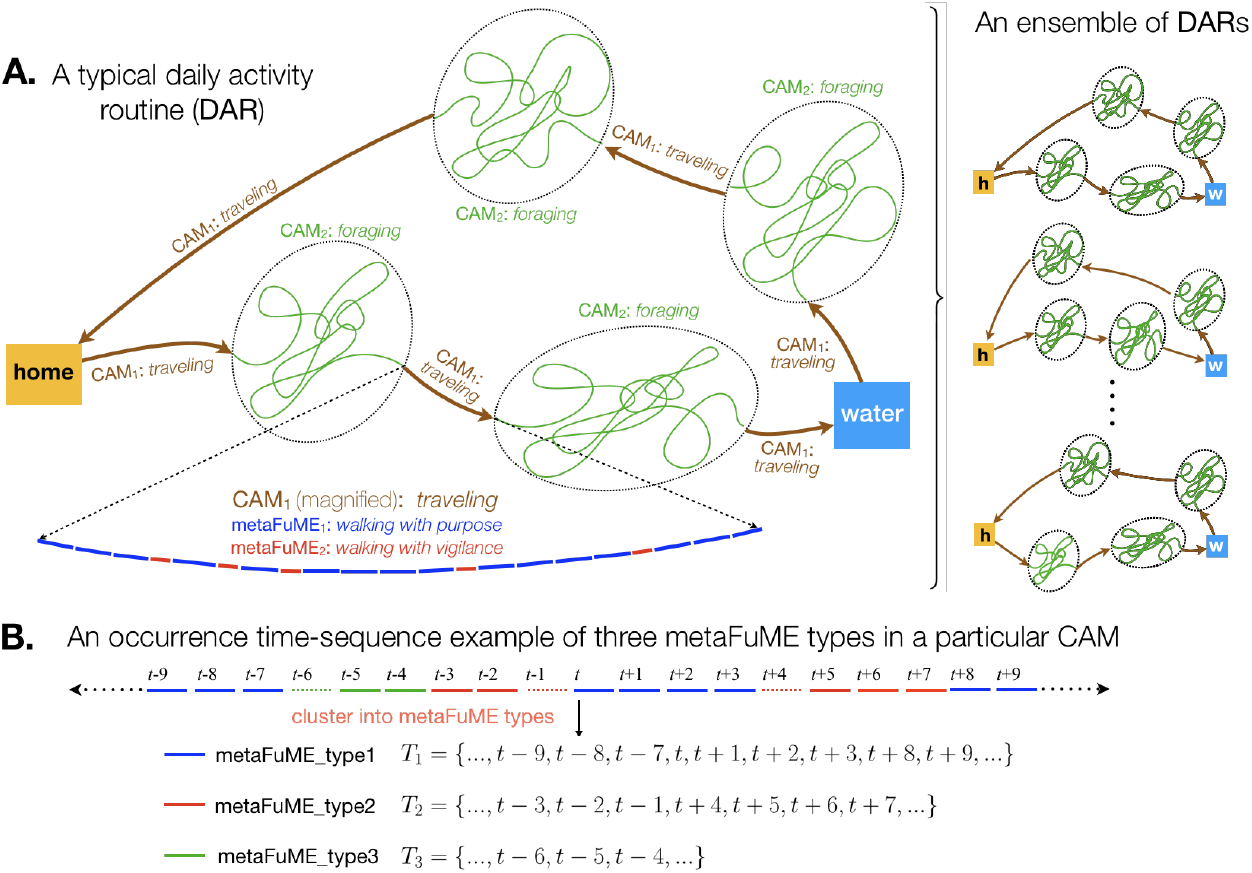
Segmentation of a daily activity routine (DAR) of a central-placed forager. **A.** Left-hand side: a simple two-CAM DAR with water stop is depicted along with a magnified depiction of a *traveling* CAM generated from two types of metaFuMEs. Right-hand side: an ensemble of DARs of a particular type (h=home, w=water). **B.** An illustration of the time sequence index sets *T_i_*, *i* = 1,2,…., obtained from an analysis of the metaFuME types that constitute a particular CAM sequence. (Figure adapted from Getz 2019)

## 2 GOALS, TASKS AND WORKFLOW

For clarity, we provide an outline of the tasks undertaken in our study and the workflow needed to carry out the required analyses and simulations. Our methods are intended to be applied to relatively high-frequency empirical data (viz., collected at frequencies around 1 to 0.01 HZ). Lacking such data, and also taking up the challenging of building models that can be fitted to such data for predictive purposes (e.g., addressing management and response to global change questions), we first focus on the building stochastic simulation models that can realistically simulate daily activity routines (DARs) with respect to their underlying CAM structure. We refer to this model as our multi-mode, biased, correlated random walk (MBCRW) model.

We use our MBCRW model to produce ensembles of DAR segments that can then be analysed as surrogates for real data. The approach we take is to extract a best-fitting basis set of metaFuMEs and associated sequence index sets that can then be used to construct a multi-CAM metaFuME Markov (M^3^) model. This M^3^ model, extracted from our surrogate data, could also be extracted directly from empirical data, collected at the appropriate frequency. When extracted either from surrogate or real data, the M^3^ model can be assessed to see how well it reproduces the CAMs evident in the original empirical/surrogate data (Fig. 2).

**Figure 2:**
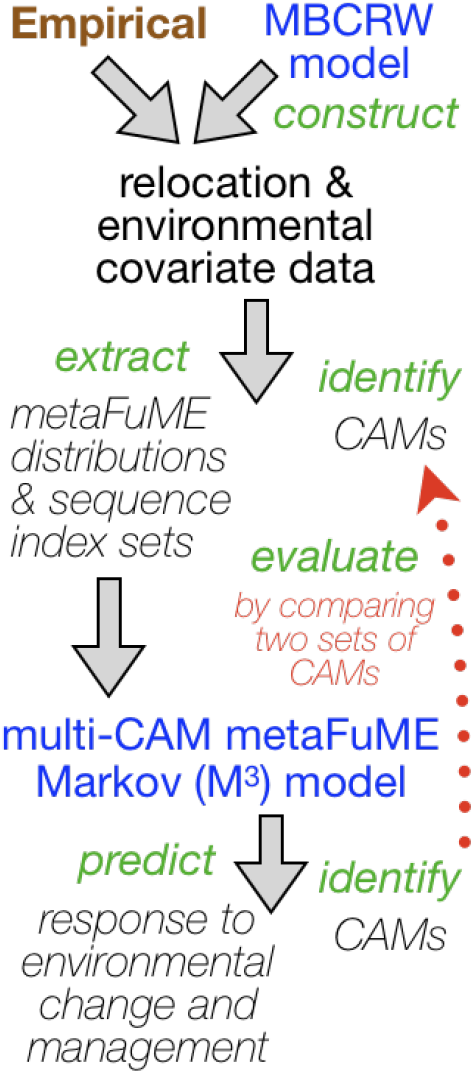
The logic behind our study. Relocation data can either be empirical or generated using a multi-mode, biased, correlated random walk (MBCRW) model. These data can then be used to extract a metaFuME distributions basis set with accompanying sequence indeces sets, as well as use various segmentation methods to identify CAMs. The metaFuME distributions and sequence index sets yield an M^3^ model that can either be used for management and prediction or evaluated in terms of how well it is able to produce the initial set of identified CAMs.

We use a stochastic process model to generate relocation data. These relocation data are created at a frequency of 0.2 Hz (i.e., locations are generated five seconds apart) for a 24 hour period (i.e., 17,280 time steps). Thus we use the model to create ensembles of DARs. The output from this model, evenly subsampled to create data sets with an order of magnitude fewer points will be used as surrogate empirical DAR segments to create ensambles 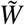(0,24*h*) (Eq. 1) relocation data sets. These data will then be used to apply our deconstructive methods and use the results to evaluate our approach. In particular, the relocation sets that we use to represent these DARs will be a subsampling of every sixth point of the generated data. This provides us with relocation sets for which one unit of *t* equals 30s, implying *t_f_* = 2,880 units over a 24 hour period. The tasks involved are the following:

A. **Simulation of DAR ensembles**

**A1.** We provide a mathematical description of the underlying simulation model as a generalized random walk with step length and directional heading correlations and biases that depend on the time of day, structure of local environment, and knowledge of the regional location of critical resources.
**A2.** We build the model using the Numerus Model Builder software platform and generate the code necessary to run our simulations.
**A3.** For a selected set of model parameter values, we run the model to obtain an ensemble of *N* DARs.
B. **Deconstruction of DAR ensembles**

**B1.** We prepare our simulated relocation data sets for analysis by thinning them from 17,280 to 2,880 points for each DAR, excluding the initial location (0,0).
**B2.** We carry out BCPA with running windows of different widths (e.g., 5, 20 and 60) or HMM analysis to identify both short and long duration CAMs.
**B3.** From the set of CAMs, we extract step length (SL) and turning angle (TA) time series and calculated means, variances, auto-correlations and covariances (cross-correlations).
**B4.** We perform cluster analysis on SL and TA point locations statistics to extract metaFuME sets and accompanying metaFuME sequence index sets.
**B5.** We group SL and TA data identified in B4 as belonging to metaFuME of a particular type, and we use the data in the sequence index sets to estimate metaFuME transition probabilities. We also estimate transition probabilities between CAMs using the CAM sequence extracted in B2.

Once A. and B. are completed, we use the metaFuME and CAM data sets and transition probabilities estimated in B5 to simulate ensembles of walks. We then compare these ensembles to those generated under Task A to assess similarities and differences. In particular, in our study we will look at SL and TA distributions, time spent in each CAM, area coverage and distance from a given point to compare the two ensambles.

## 3 MODEL: DESCRIPTION AND IMPLEMENTATION

In terms of movement models, two contrasting approaches are common. The first is the use of numerical event-oriented (e.g. Gillespie’s event oriented algorithm— (Gillespie 1976, Gillespie 1977), or time-discretizations methods to compute solutions to stochastic differential equations that model continuous time stochastic processes. In increasing order of complexity, these include a pure random walk (Wiener process; e.g. see (McClintock et al. 2014)), a linear- and angular-velocity-biased random walk (Ornstein-Uhlenbeck process, (Gurarie et al. 2017)), and location salient random walk (e.g., walks in a force field, (Magdziarz et al. 2012); or movement dependent on the local landscape, (Harris and Blackwell 2013)), where the location itself could be a function of time (e.g., the centroid of a moving group—see (Langrock et al. 2014)). The second is to use rule-based simulations of movement over rectangularly or hexangonally rasterized landscapes without the benefit of a compact underlying differential equation structure (Getz et al. 2015, del Mar Delgado et al. 2018)

Our path simulation algorithm of the movement of individuals over landscapes is an outgrowth of Brownian motion (Pozdnyakov et al. 2014), correlated random walks (Kareiva and Shigesada 1983) and, even Levy walks (Benhamou 2007) simulation methods. Within the context of appropriate modeling methods (Getz et al. 2018), our elaborations to these earlier approaches in formulating our MBCRW include both biased directionality and mixed-distribution (Morales et al. 2004) components. The multiple modality aspect of our model arises in two ways. First, from switching movement modes locally at a frequency that assumes a particular level of landscape heterogeneity (e.g., moving within and among resource acquisition patches). Second, from switching directional movement biases at some slower daily-activity-related frequency in terms of attraction to distant (peripheral) geographic locations such as water holes or resource rich areas, and then attraction back to centrally located nest sites or home range centers. Detailed mathematical description of this model can be found in the supporting online file (SOF). In short, our model is an MBCRW, with daily walks between and around two points of attraction, that always starts around the same central point, followed by a daily excursion towards and around a peripheral point that may change each day, with a final end-of-each-day movement back towards the central point (see right-hand panel in Fig. 3).

**Figure 3:**
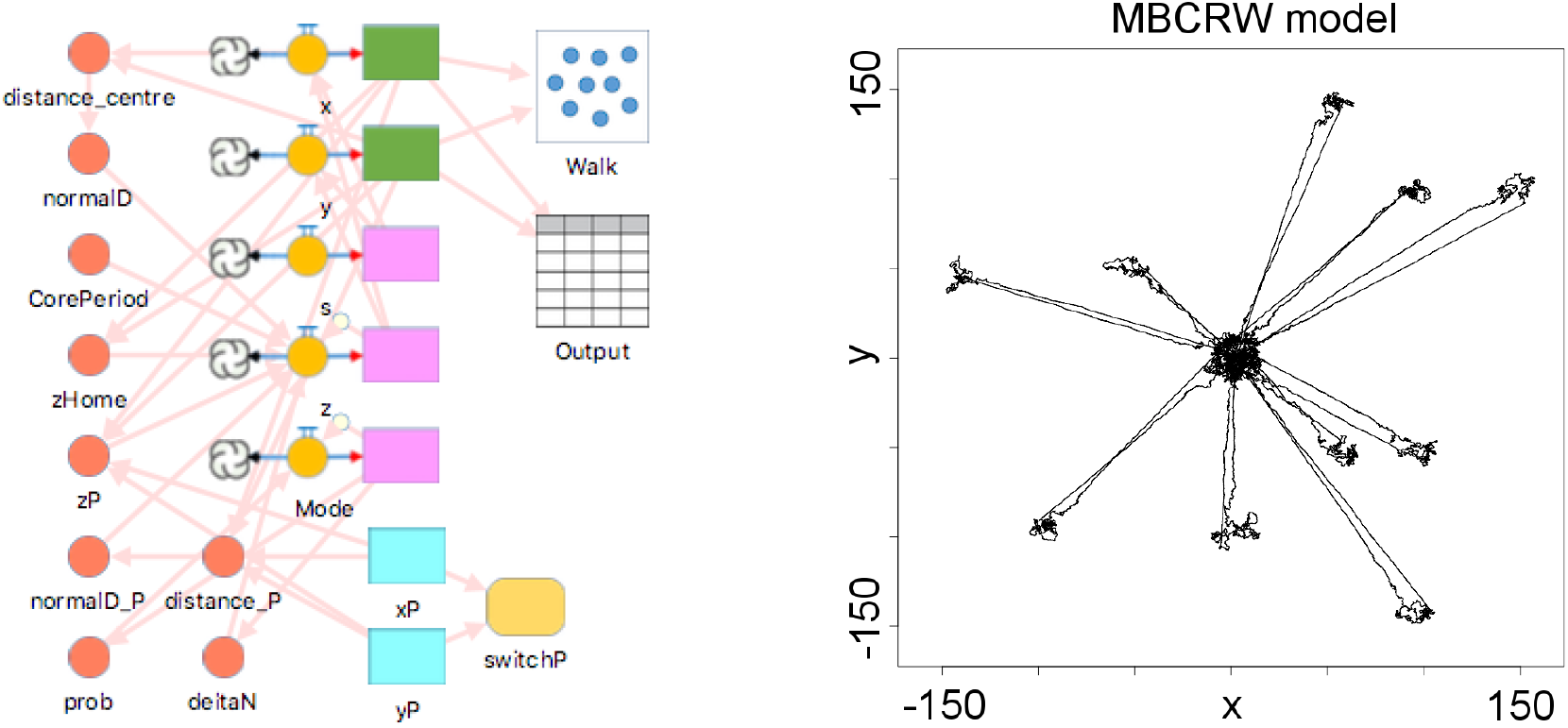
A graphical construction of our NMB model (left panel; values and equations are inserted into a form that is opened when a component icon is selected) and output from one instance of a stochastic 10-day simulation (right panel)

Our model is implemented using the Numerus Model Builder (NMB) platform, which allows us to code a digital simulation using a set of graphically implemented, drag-and-drop icons, as shown in Figure 3, in which mathematical expressions of model equations are inserted. Once all the necessary equations and segments of code have been inserted into equation windows and code chip frames (Getz et al. 2015, Getz et al. 2018), NMB then generates a script that can be used to run simulations of the model over the desired interval of time. The NMB modelling panel and the output of the simulation for the MBCRW model are shown in Fig. 3. We generate the set of relocation data 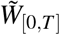 by simulating the model for a 10-day temporal horizon (*T* = 172,800), where one time-unit in NMB corresponds to 5s and one spatial unit to 100m. In SOF we provide the parameters used in this simulation.

## 4 DECONSTRUCTION OF DAR ENSEMBLES

The approach we develop here is to use the time interval that corresponds to the same frequency at which the data were collected and to either use actual step-size and turning-angle data, or smoothed/idealized functions that the are fitted to or theoretically represent these data. As part of the simulation, considerations of directional biases, step-size and turning-angle serial correlations, step-size/turning-angle cross-correlations, and context-dependent environmental and individual internal-state step-size and turning-angle distributions may be considered (Ahearn et al. 2017, Langrock et al. 2014). Although, individuals may also take the locations and directions of heading of other individuals into account as they move across landscapes (Couzin et al. 2005, Conradt et al. 2009, Langrock et al. 2014), we do not consider this level of complexity here.

We undertook analyses of the relocation data using the R Studio implementation of the R programming language, where in our case these data were obtained in a CSV format from our NMB simulations. Our analyses were designed to extract from the data all the information needed to construct our M^3^ model.

First, we subsampled the output of the NMB simulation data at a frequency appropriate for extracting relatively fast canonical activity modes (i.e., CAMs at a sub-hourly to hourly scale—see Getz 2019) by selecting every 60^th^ point, which corresponds to a 5 min interval between relocation points. Using the R package `moveHMM`, we then applied the Viterbi algorithm to sort our subsampled relocation data into the most likely sequence of CAMs and identified every distinct CAM as one of *K* types. Each CAM segment can be represented by

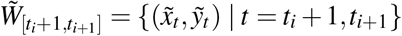

such that a complete path 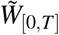 is then made up of *p* such segments strung together over an interval [0, *T*]: i.e.,

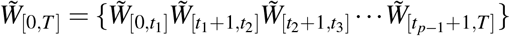

with a corresponding set of *p* – 1 change points *T*_CP_ = {*t*_1_, *t*_2_, …, *t*_*p*–1_}.

Second, we subsampled the output of the NMB simulation data at a frequency appropriate for extracting metaFuMEs (i.e., around 10 s to 1 min—see Getz 2019) by selecting every *5^th^* point, which corresponds to a 30 s interval between relocation points. Thus, given the extracted sequence of CAM at a 5-min relocation point interval, we assigned the same CAM index to each of the corresponding 10 time-points included in each step of the 5-min interval identified in the first step as belonging to a particular CAM type.

For each resulting segment at a 30 s time-scale, we extracted the step length *s*(*t*) and turning angle *a*(*t*) time-series, by calculating the following values:

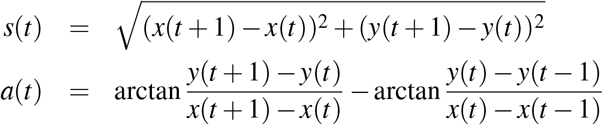

for *t* = 1,…,*T*. Note that for *t* = 0 both values are not defined. We then obtained the mean and the variance (*μ_x_, σ_x_*), where *x* = *s, a*, and calculated the running-term auto-correlation timeseries *v^xx^*(*t*) for *t* = 2,…,*T* and running-term cross-correlation time-series *v^as^*(*t*) for *t* = 1,…,*T*:

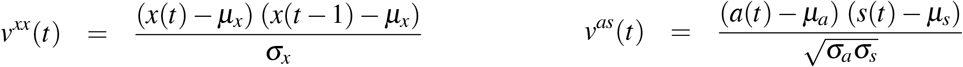

We also computed the time-series of the absolute value of the turning-angle cosines, which we subsequently used in our metaFuME cluster analysis in place of the the turning angles themselves: i.e., we generated the values

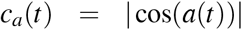

We performed data standardization through min-max normalization (Van Moorter, Visscher, Jerde, Frair, and Merrill 2010), as data pre-processing before carrying out a cluster analysis:

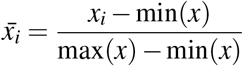

We collected the sets 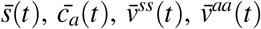 and 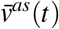 for each *t* identified as part of the same CAM to perform cluster analysis. We used Ward’s method, provided by the R package `stats`, to extract the set of clusters of metaFuMEs for each CAM and the elbow method to estimate the optimal number of clusters.

Once we have extracted the clusters of metaFuMEs for each CAM, we evaluated if some metaFuME clusters could be associated and grouped together. To accomplish this, we performed cluster analysis of mean and variance of SL and TA values for each metaFuME initial cluster, again using Ward’s method and then estimating the optimal number of clusters through the elbow method. The results of the elbow method are shown in the SOF (Figure S2).

Once we have identified the final clusters of *J* metaFuMEs, we extract the corresponding sets of step-lengths *S_j_* and turning-angles *A_j_* for each metaFuME type *j* = 1,…, **J**. In our study, when simulating the M^3^ model, we sampled from these data, according to the chosen or current metaFuME type such that each element in *S_j_* and *A_j_* had an equal probability of being selected.

As a following step in our analysis, we extracted the sequences of metaFuME types for each CAM segment. Using the R package `markovchain`, we fitted the transition probability matrix for each sequence and calculated the weighted average (depending on the sequence length) to extract the metaFuME transition probability matrices for each CAM. Given *n* sequences for a given CAM, the weighted average for a matrix entry was calculated as follows:

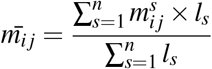

where *l_s_* is the length of the sequence *s* and 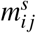 is the (*i, j*) entry of the matrix extracted from sequence *s*. Note that some CAMs might not contain all the *J* metaFuMEs. Assuming that *M_s_k__* indicates the index set of the metaFuME types that compose a given CAM *k*, for *i* ∉ *M_s_k__* the matrix *M_k_* will need to contain entries *m_ij_* equal to 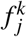, for *j* ∈ *M_s_k__*, and equal to 0, for *j* ∉ *M_s_k__*, where 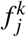 represents the frequencies of metaFuME *j* in CAM *k*. Moreover, the entries *m_ij_*, for all *i* and *j* ∉ *M_s_k__*, will be equal to 0. Defining these rows in the transition probability matrices, facilitates the change to the correct metaFuME sets when switching between CAMs.

Recalling the first task of our analysis, we identify the most likely sequence of CAMs again using the Viterbi algorithm. In this case, however, we used the R package `markovchain` and the CAM sequence to extract the CAM transition probability matrix *M_C_*. In our simulation, we assumed the same CAM always persisted for at least 10 time points (i.e., 5 mins) and therefore we evaluated the transitions between CAMs using *M_C_* every 10 time points.

## 5 M^3^ MODEL SIMULATION

By performing the tasks defined in Section 2, and described in more detail in Section 4, we extracted all the data related to metaFuMEs and CAMs which are necessary to construct an M^3^ model. To summarise, these data are the following: a finite set of *J* metaFuME SL and TA variables set *S_j_* and *A_j_*, a *K × K* transition probability matrix *M_C_*, representing the probability to switch between CAMs, and *K, J × J* transition probability matrices *M_k_, k* = 1,…, *K* for each of the *K* CAM types, to describe the transition probabilities between metaFuMEs while in a given CAM type.

The simulation of this model produces a set of relocation data 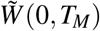, as described in Algorithm 1. This algorithm is an amalgam of an “outer” and “inner” process. The outer process is a Markov transition process governing the probability that the current CAM will switch to a different CAM at every 5-min time interval. The inner process is a within-CAM stochastic dynamical systems process that controls the change of location from one time-step to the next (30 s time-scale) according to the selected metaFuME.

### Algorithm

To initialize the simulation, we declare the initial location (*x*_0_, *y*_0_), initial CAM *k*, initial metaFuME *j*, initial heading *θ*_0_ and final time *T_M_*. For each time step *t*, we generate location data by drawing step length and turning angle (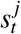 and 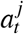) pairs from the sets *S_j_* and *A_j_*, according to the metaFuME type *j* in operation at time *t*. The location data and the elapsed times are updated every step and the switch between metaFuME types is ruled by the matrix *M_k_* (when in CAM *k*). After the given time in CAM *k* has elapsed (10 time steps, counted using the variable 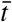), the CAM update is defined by the transition probability matrix *M_c_*. We use the function *f* to represent the process by which both metaFuMe and CAM values are updated according to their current values and the given transition probability matrices.

#### Algorithm 1: M^3^ model simulation

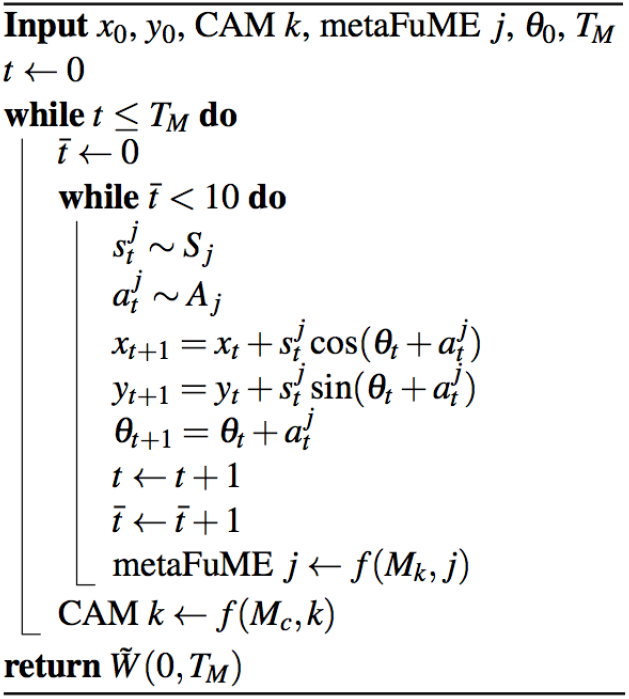

## 6 M^3^ MODEL PARAMETERS

In our study, the extracted M^3^ model has the following features. It is composed of 3 CAM types and 3 metaFuME types. Results of the HMM analysis are shown in Fig. 4.

**Figure 4:**
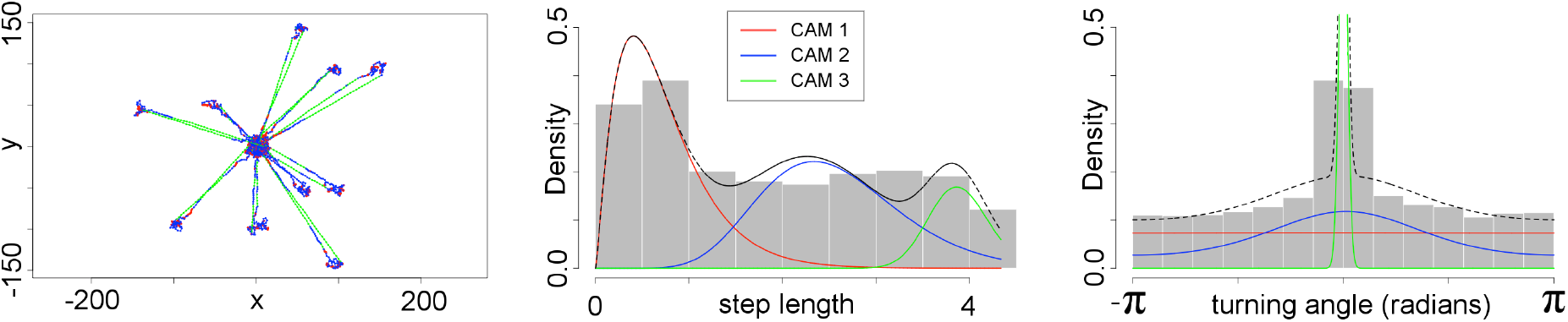
Results of the CAM extraction from our HMM analysis (plotted using the R package moveHMM). Left panel: CAM segments plotted using legend colors. Middle panel: SL distribution in 100m bins. Right panel: TA distribution in radians.

From Fig. 4, we see that CAM 1 represents random movement mostly around the centers of attractions while CAM 2 represents more correlated movement around these centers. On the other hand, CAM 3 represents highly directed movement that occurs between centers of attraction. From our cluster analysis (details in SOF) CAM 1 is composed of metaFuMEs of type 1 and 2 (frequencies: 0.51, 0.49), CAM 2 of metaFuMEs of type 1, 2 and 3 (frequencies: 0.4, 0.1, 0.5) while CAM 3 by type 1 and 3 (frequencies: 0.16, 0.84). Distributions of SL and TA for the 3 metaFuME types are shown in Fig. 5. MetaFuME of type 1 is a *variable step/persistent direction* metaFuME while metaFuME of type 2 is a *variable step/random direction* one. MetaFuMe 3 is a *large step/persistent direction* metaFume type.

**Figure 5:**
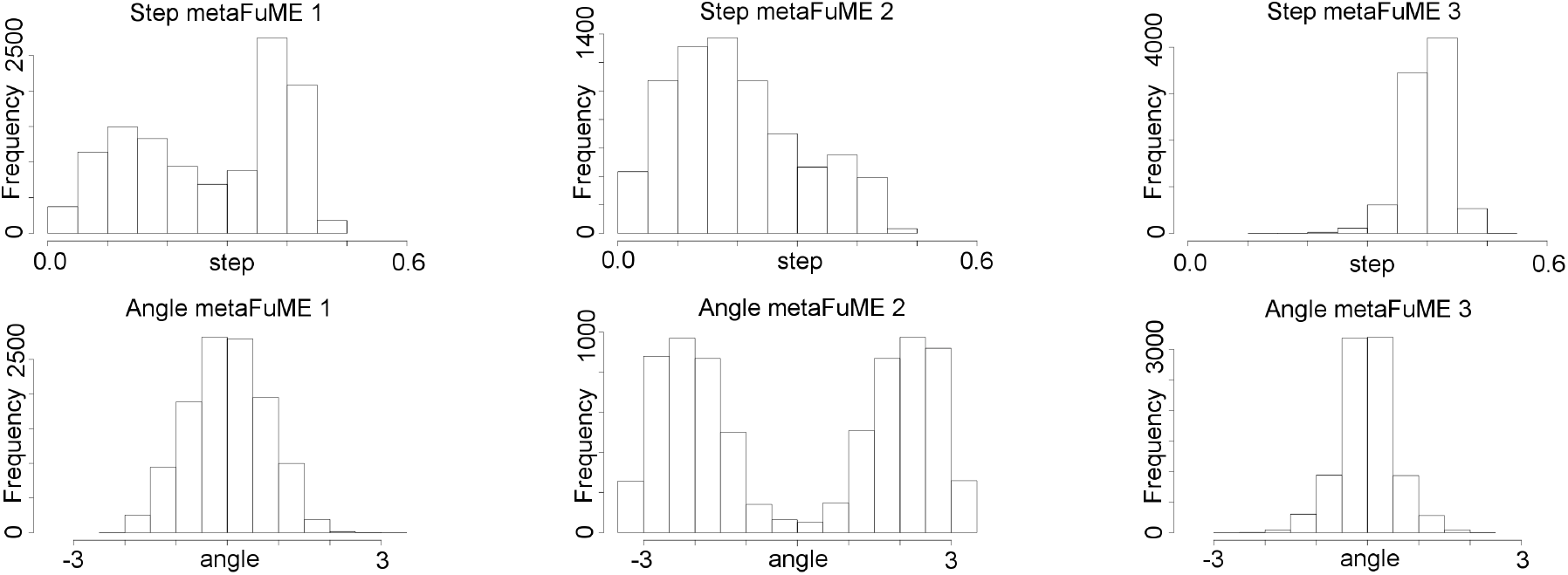
Distribution of SL (100m bins) and TA (radians) for metaFuME types, from subsampled MBCRW data

The CAM transition probability matrix *M_C_* and the matrices *M_k_* for each CAM of type *k* = 1,…, *K* for the M3 analysis of the MBCRW data generated for this study can be found in our SOF. The initial configuration of our M^3^ model was (0, 0) for the initial location, CAM 1 as initial CAM and metaFuME 1 as initial metaFuME. We implemented this model in NBM and run a simulation for 28,800 time steps (10 days). The NMB modelling panel and the output of the simulation are shown in Fig. 6. Additional examples of the M^3^ model simulation output are provided in our SOF (Figure S3).

**Figure 6:**
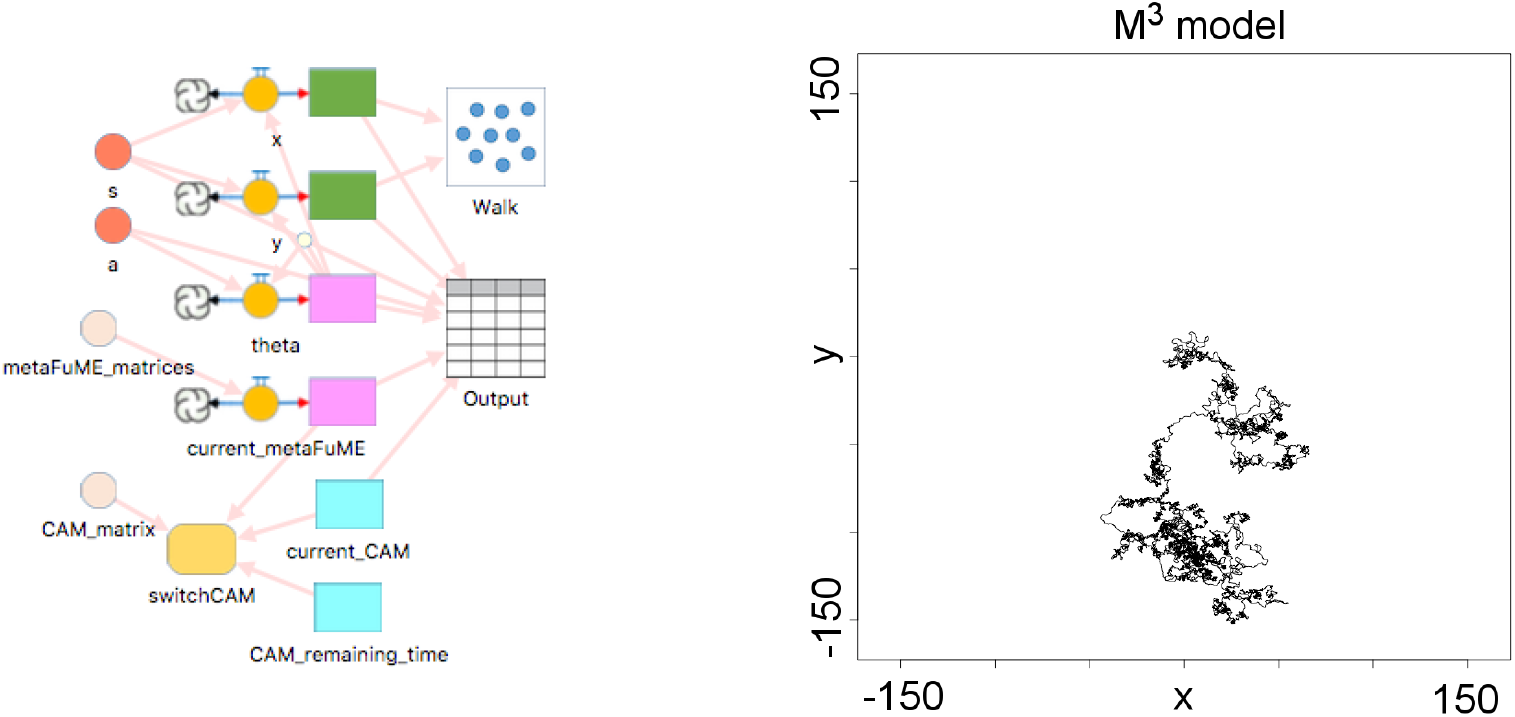
NMB model canvas (left panel) and one instance of a simulation (right panel) of our M^3^ model

## 7 MODEL COMPARISONS

The structure of the paths obtained from of our MBCRW and M^3^ models are very similar at the metaFuME and multi-CAM scales by design. This is illustrated in SOF where the SL and TA distributions are plotted side by side in Fig. S4 and S5 for each of the CAM types. We used the R package `trajr` to calculate the sinuosity of the two simulated paths and, unsurprisingly, obtained similar results because sinuosity is a relatively fine scale structure of a movement path: viz., 2.75 for the MBCRW model and 2.90 for the M^3^ model. Moreover, we extracted the time spent in each CAM, for both models (Fig. S6 of SOF) and these were similar as well. The two models, as expected, varied quite considerably, when comparing the more global scale features of the paths they produce. In particular, the total area covered over time (Fig. S7 in SOF), the area covered each day (Fig. S8 in SOF) and the distance from the origin over time (Fig. S9 in SOF) were notably different. These results confirm that the M^3^ model captures local features of movement paths (e.g., SL and TA distributions and sinuosity) but fails to capture capture global features which arise in our MCBRW model from movement bias due to the presence of centers of attraction that periodically switch over time. Thus, derived statistical movement models, such as M^3^, need to be improved to capture global features by considering not only turning angles, but actual angles of heading.

## 8 CONCLUSION

Recently, stochastic walk models that do not include switches in directional movement bias beyond metaFuME and CAM time scales, have been promoted as a way to capture global features of walks from empirical relocation data—e.g., the use of autocorrelated-kernel-density estimation (AKDE) to compute home range size (Fleming and Calabrese 2017, Noonan et al. 2019) from movement paths. Such methods, however, as demonstrated through our comparison of output from our MBCRW and M^3^ models, will fail to provide reliable estimates of global properties of walks when the relocation data is fitted to models that fail to take directional biases into account at the multi-CAM scale. This includes the Ornstein-Uhlenbeck stochastic process model that is the bases for path analysis and home range estimation using the increasingly popular `ctmm` R package (Calabrese, Fleming, and Gurarie 2016). As with the Ornstein-Uhlenbeck model, our M^3^ models fails to take key processes that primarily affect global rather than local structure of paths and thus needs to be extended to take such processes into account. The answer is not to try to fit our MBCRW model directly to relocation data since, as with any approach that tries to simultaneously fit data created by processes occurring at different scales, the approach is likely to dramatically fail. Rather, an extended M^3^ that first fits an M^3^ model to relocation data, as outlined here, and then accounts for processes taking place at the multi-CAM time scale and beyond, is needed; and is a topic that remains to be addressed in future studies.

## Supporting information

SOF

## Acknowledgments and SOI File

The development of NOVA, a precursor to Numerus Model Builder, was supported by NSF grant CNS-0939153 to Oberlin College (PI: RS) and NSF-EEID grant 1617982 (PI: WMG). The supporting online file (SOF), containing details of the MBCRW model, simulation parameter values, and additional results is available at https://ludovicalv.github.io/PDFs/SOF_SCS_2020.pdf.

## AUTHOR BIOGRAPHIES

**WAYNE M. GETZ** is a Professor in the Graduate Division at the University of California, Berkeley, and a Founding Partner of Numerus Inc. His email address is.

**LUDOVICA LUISA VISSAT** is a Postdoctoral Scholar at the Department of Environmental Science, Policy and Management at the University of California, Berkeley. Her email is.

**RICHARD SALTER** is an Emeritus Professor of Computer Science at Oberlin College and a Founding Partner of Numerus Inc. His email address is.

