## Supplementary material for "Simulation and Analysis of Animal Movement Paths using Numerus Model Builder": SOF

### 1 MATHEMATICAL DESCRIPTION: MBCRW MODEL

We now introduce the mathematical description of the MBCRW model. This model describes a MBCRW with alternating centres of attraction $O$ and $P_i$. Each daily movement will involve a different $P_i$. The centre of attraction, indicated with $P$, switches as follows:

$$P = \begin{cases} O & \text{if } cos(\frac{2\pi t}{T}) > 0 \\ P_i & \text{otherwise on day } i \end{cases}$$

where $t$ indicates the elapsed time and $T$ represents the length of the day. At time $t = 0$ $P$ will be equal to $O$, switching after $\frac{T}{4}$ to $P_i$ and to $O$ again after $\frac{3}{4}T$.

**Location data.** Given the initial position $(x_0, y_0)$, the location coordinates are updated as follows:

$$\begin{aligned} x_{t+1} &= x_t + s_t \cos\theta_t \\ y_{t+1} &= y_t + s_t \sin\theta_t \end{aligned}$$

where $s_t$ is the step length and $\theta_t$ is the absolute heading. Note that an equivalent description could define $\theta_{t+1}$ as function of the turning angle $\alpha_t$, as follows:

$$\theta_{t+1} = \theta_t + \alpha_t$$

and apply the trigonometric functions to $(\theta_t + \alpha_t)$.

**Step-length.** $s_t$ is drawn from the uniform distribution:

$$s_t \sim \text{UNIFORM}(s_{min}, s_{max})$$

**Absolute heading.** $\theta_t$ varies with time and location. We define $\theta_t$ as follows, where $P$ represents the centre of attraction at time $t$:

$$\theta_{t+1} = \begin{cases} \theta^{c_P} & \text{if } z^u > f(d_t^P) \\ \theta_t + \theta^n & \text{otherwise} \end{cases}$$

$\theta^{c_P}$ is drawn from the distribution:

$$\theta^{c_P} \sim \text{UNIFORM}(\theta^P - (1 - corrP) \times \frac{\pi}{2}, \theta^P + (1 - corrP) \times \frac{\pi}{2})$$

where $\theta^P$ is the absolute heading towards the point $P$ (function of the animal current position) and $corrP$ defines the heading correlation (0: no correlation, 1: maximum correlation).

$z^u$ is drawn from the uniform distribution:

$$z^u \sim \text{UNIFORM}(0,1)$$

while the function $f$ is defined as follows:

$$f(d_t^P) = \frac{1}{1 + (d_t^P / d^{\text{crit}_P})^{\eta_P}}$$

This function controls the location of attraction and therefore the range of movement. The parameter $d_t^P$ is the distance from point P at time $t$, with $d^{\text{crit}_P} > 0$ and $\eta_P > 1$.

The random variable $\theta^n$ is drawn from the distribution:

$$\theta_n \sim \begin{cases} \text{UNIFORM}(-\pi, \pi) & \text{if } m = 1 \\ \text{UNIFORM}(-(1-\text{corr}H) \times \pi, (1-\text{corr}H) \times \pi) & \text{if } m = 2 \end{cases}$$

with heading correlation corr$H$. $m$ describes 2 different modes: the first mode represents an uncorrelated random walk, within the critical distance, while the second one represents a walk with heading correlation greater than 0. The value of $m$ can switch between the 2 values accordingly to the following transition matrix:

$$\begin{pmatrix} \theta_{11} & 1-\theta_{11} \\ 1-\theta_{22} & \theta_{22} \end{pmatrix}$$

with $\theta_{11}, \theta_{22} \in [0,1]$.

### 2 SIMULATION PARAMETERS

To define the parameters of our model, we established 5s as a time unit (17,280 in one day) and one space unit equal to 100m. In 5s, the travelled distance is around 7m (assuming straight walk and speed $\approx$ 5 km/h). We run the simulation for 172,800 time units (10 days); we set as initial location the origin O: (0,0), initial mode $m = 1$ and the centres of attraction $P_i$ as shown in Figure S1. These points have the following coordinates: (-150,50), (-100,-100), (-50,50), (0,-100), (50,-50), (50,150), (100,-150), (100,-50), (100,100) and (150,100).

We use the following set of parameters:

| Param name | Value | Param name | Value |
|---|---|---|---|
| $s_{min}$ | 0.05 | $\eta_P$ | 3 |
| $s_{max}$ | 0.09 | corrH | 0.8 |
| $T$ | 17280 | corrO | 0.8 |
| $d^{crit_O}$ | 50 | corrP$_i$ | 0.8 |
| $d^{crit_{P_i}}$ | 50 | $\theta_{11}$ | 0.995 |
| $\eta_O$ | 3 | $\theta_{22}$ | 0.995 |

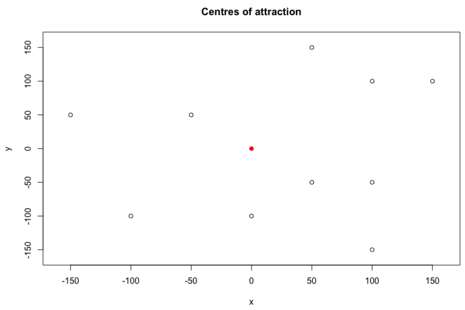


Figure S1: Centers of attraction $O$ (in red) and the other 10 centers $P_i$. Note that the scale represents the spatial units in NMB.

### 3 CLUSTERING ANALYSIS

The elbow method results are shown in Figure S2. We used this method to choose the optimal number of clusters for the metaFuMe clusters, for each CAM and in total.

In the following table we report the number of times the different metaFuMes types occurred in each CAM, from the analysis of the subsampled data of the MBCRW model.

| CAM | metaFuME 1 | metaFuME 2 | metaFuME 3 |
|---|---|---|---|
| 1 | 6695 | 6314 | 0 |
| 2 | 4560 | 1093 | 5715 |
| 3 | 612 | 0 | 3205 |

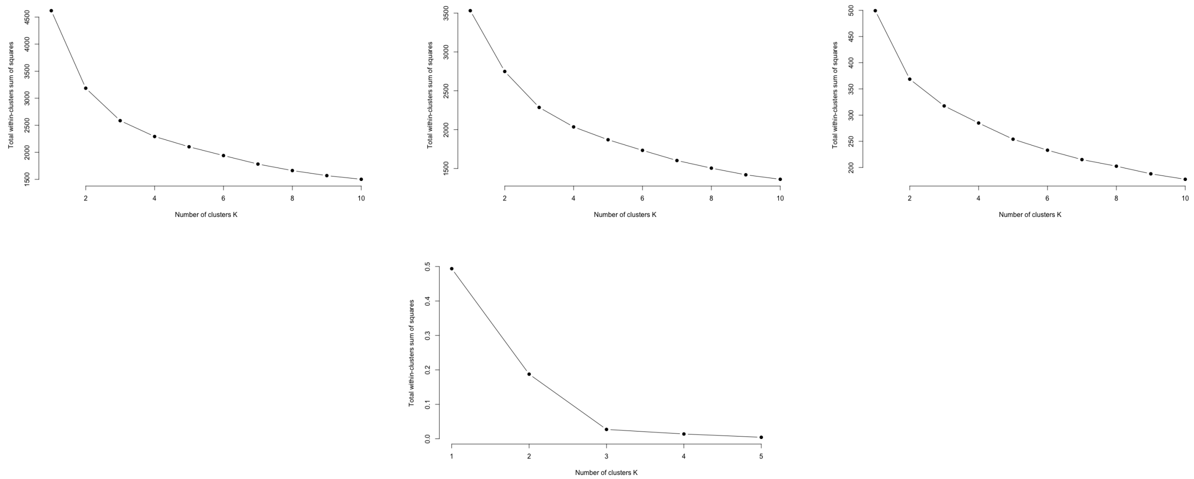


Figure S2: Elbow method - for each CAM (first row) and total (second row)

### 4 M$^3$ PARAMETERS

$$M_C = \begin{bmatrix} 0.80 & 0.20 & 0.00 \\ 0.22 & 0.75 & 0.03 \\ 0.00 & 0.09 & 0.91 \end{bmatrix}$$

$$M_1 = \begin{bmatrix} 0.54 & 0.46 & 0 \\ 0.48 & 0.52 & 0 \\ 0.51 & 0.49 & 0 \end{bmatrix} \qquad M_2 = \begin{bmatrix} 0.47 & 0.27 & 0.26 \\ 0.39 & 0.24 & 0.37 \\ 0.38 & 0.05 & 0.57 \end{bmatrix} \qquad M_3 = \begin{bmatrix} 0.43 & 0 & 0.57 \\ 0.16 & 0 & 0.84 \\ 0.13 & 0 & 0.87 \end{bmatrix}$$

### 5 EXAMPLES OF M$^3$ MODEL SIMULATIONS

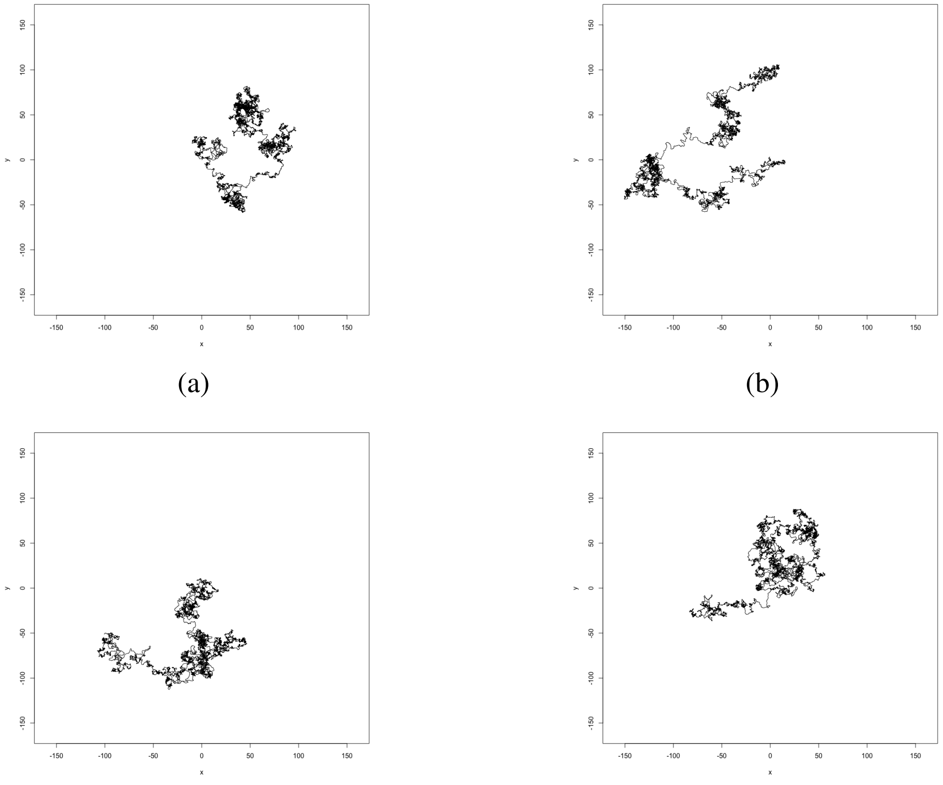

(a) (b) (c) (d)

Figure S3: Four examples of M$^3$ model simulations ($T$ = 28,800) with sinuosity values: (a) 2.975889, (b) 2.88076, (c) 2.943064, (d) 2.910309

### 6 MODEL COMPARISON

Step-length distribution for the MBCRW and M$^3$ model are shown in Figure S4, while turning-angle distributions for both models are shown in Figure S5. In Figure S6 we show the distribution of time spent in each CAM for both models. The total area covered and the area covered each day are shown in Figures S7 and S8, while the distance from the origin over time is shown in Figure S9, for both models.

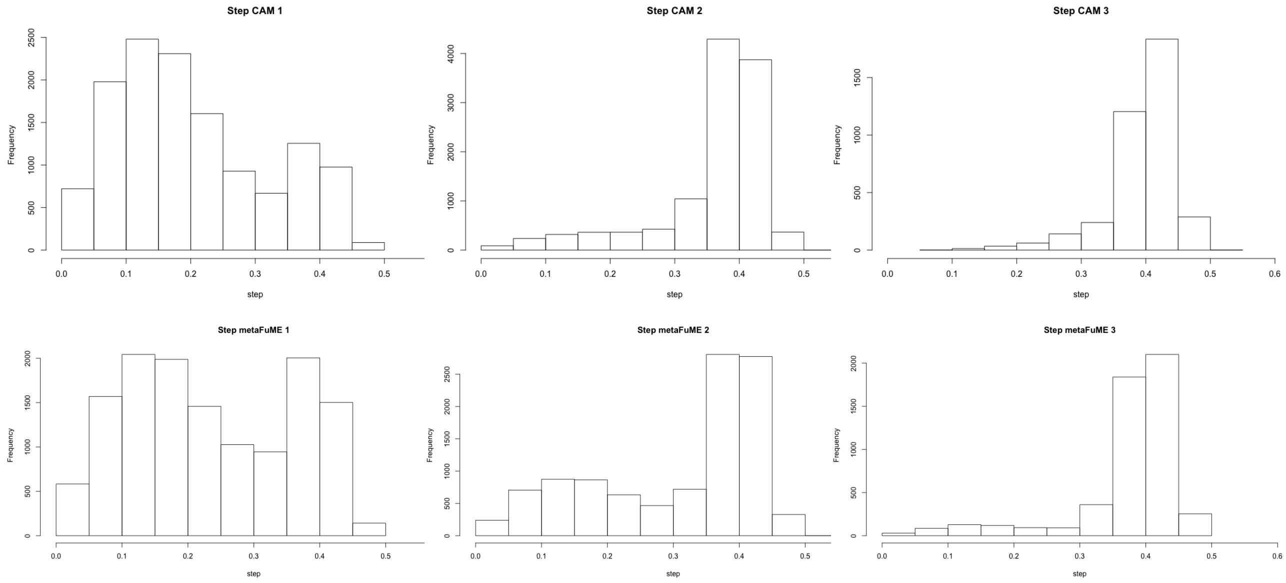


Figure S4: Step-length distribution for the different CAMs: MBCRW model (first row) and $M^3$ model (second row)

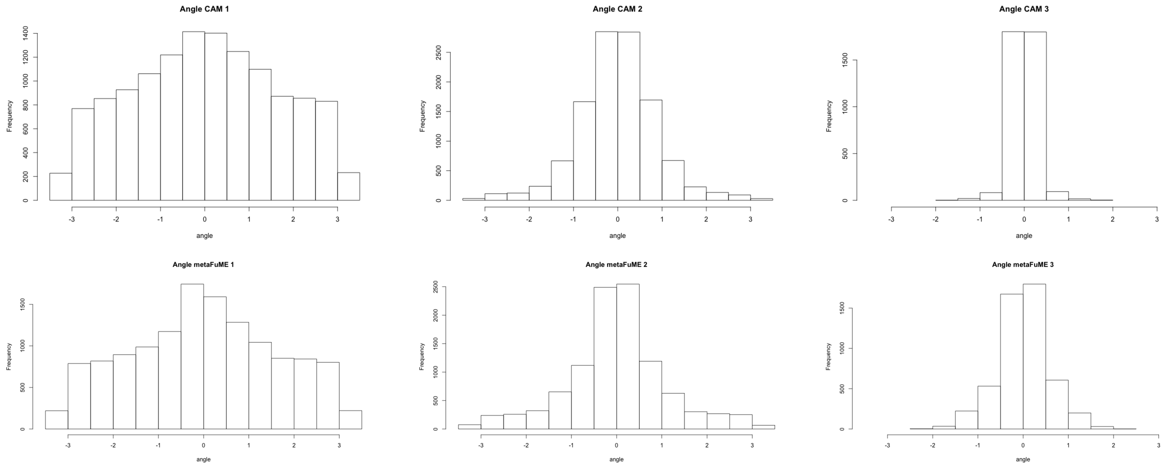


Figure S5: Turning-angle distribution for the different CAMs: MBCRW model (first row) and $M^3$ model (second row)

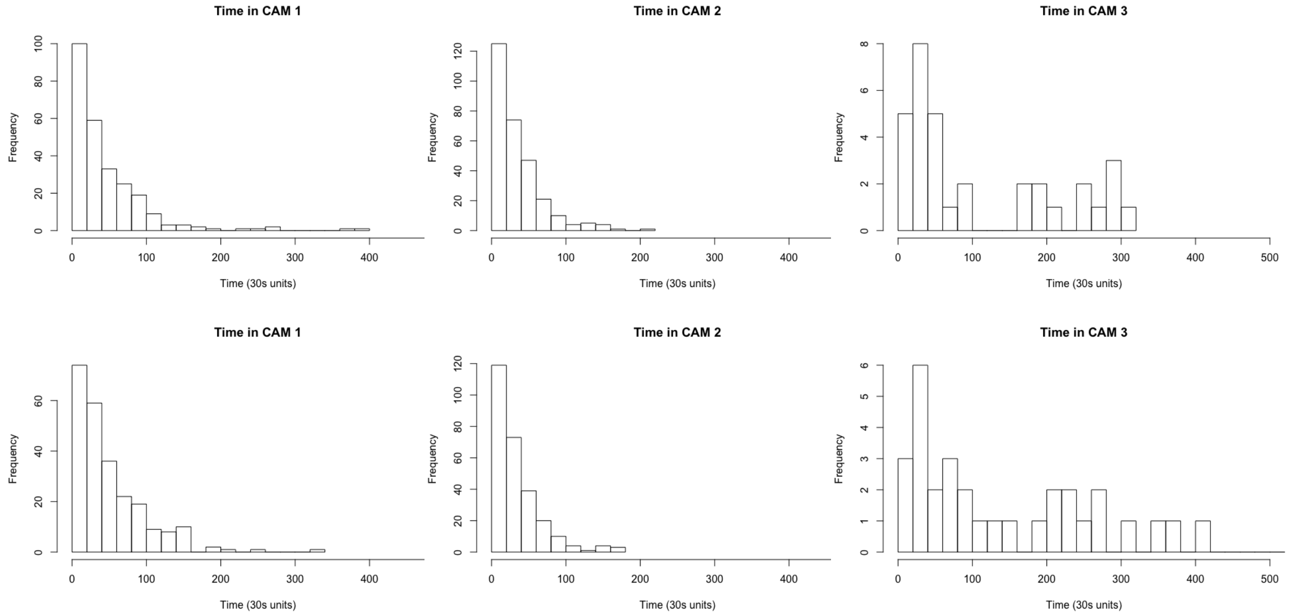


Figure S6: Distribution of time spent in the different CAMs: MBCRW model (first row) and $M^3$ model (second row)

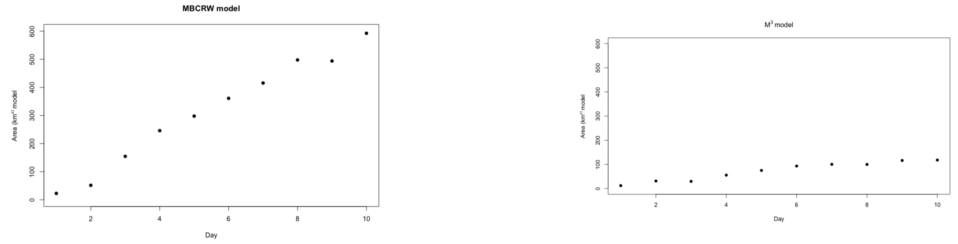


Figure S7: Total area coverage over time for the two models: MBCRW on the left, $M^3$ on the right.

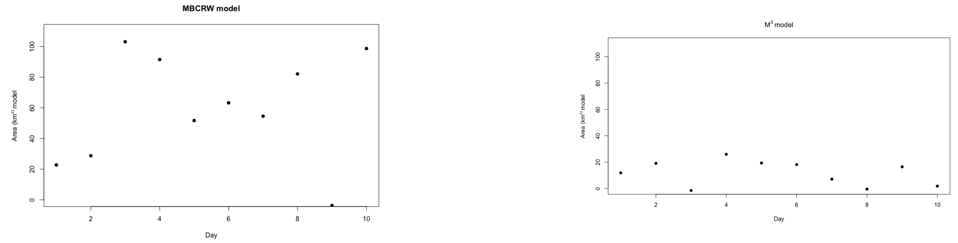


Figure S8: Area coverage each day for the two models: MBCRW on the left, $M^3$ on the right.

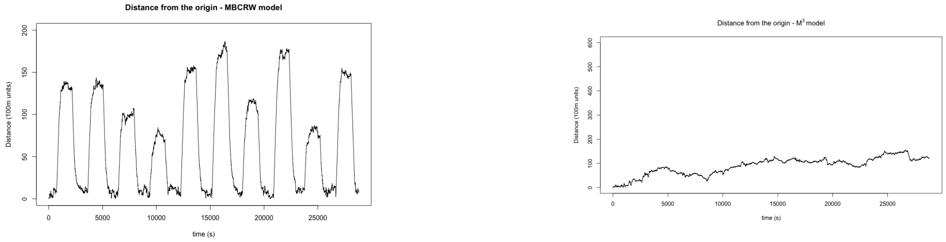


Figure S9: Distance from the origin over time for the two models: MBCRW on the left, $M^3$ on the right.
